# The Brain Computes Dynamic Facial Movements for Emotion Categorization Using a Third Pathway

**DOI:** 10.1101/2024.05.06.592699

**Authors:** Yuening Yan, Jiayu Zhan, Oliver G. Garrod, Chaona Chen, Robin A.A. Ince, Rachael E. Jack, Philippe G. Schyns

## Abstract

Recent theories suggest a new brain pathway dedicated to processing social movement is involved in understanding emotions from biological motion, beyond the well-known ventral and dorsal pathways. However, how this social pathway functions as a network that computes dynamic biological motion signals for perceptual behavior is unchartered. Here, we used a generative model of important facial movements that participants (N = 10) categorized as “happy,” “surprise,” “fear,” “anger,” “disgust,” “sad” while we recorded their MEG brain responses. Using new representational interaction measures (between facial features, MEG_t_ source, and behavioral responses), we reveal per participant a functional social pathway extending from occipital cortex to superior temporal gyrus. Its MEG sources selectively represent, communicate and compose facial movements to disambiguate emotion categorization behavior, while occipital cortex swiftly filters out task-irrelevant identity-defining face shape features. Our findings reveal *how* social pathway selectively computes complex dynamic social signals to categorize emotions in individual participants.

## Introduction

As a social species, humans possess a remarkable tool for communication^1^–the face–which enables senders to convey abundant social information^2,3^, which is efficiently and accurately decoded by the receiver’s brain (Figure 1). In such interactions, static facial features such as shape and complexion convey stable traits like identity^4,5^, age^6,7^, sex^8,9^, ethnicity^10,11^ and attractiveness^12^. In parallel, the face can also convey transient states, such as emotions^13–16^, attitudes^17–20^ and intentions^21,22^, via its dynamic facial movements–called Action Units (AUs) in the Facial Action Coding System^23^. Orchestrated by independent striated muscles, these movement features can be activated individually (e.g., lip corner puller) or collectively (e.g., smiling involving lip corner puller, cheek raiser and dimpler) to produce facial expressions of emotions^1,23,24^.

**Figure 1.**
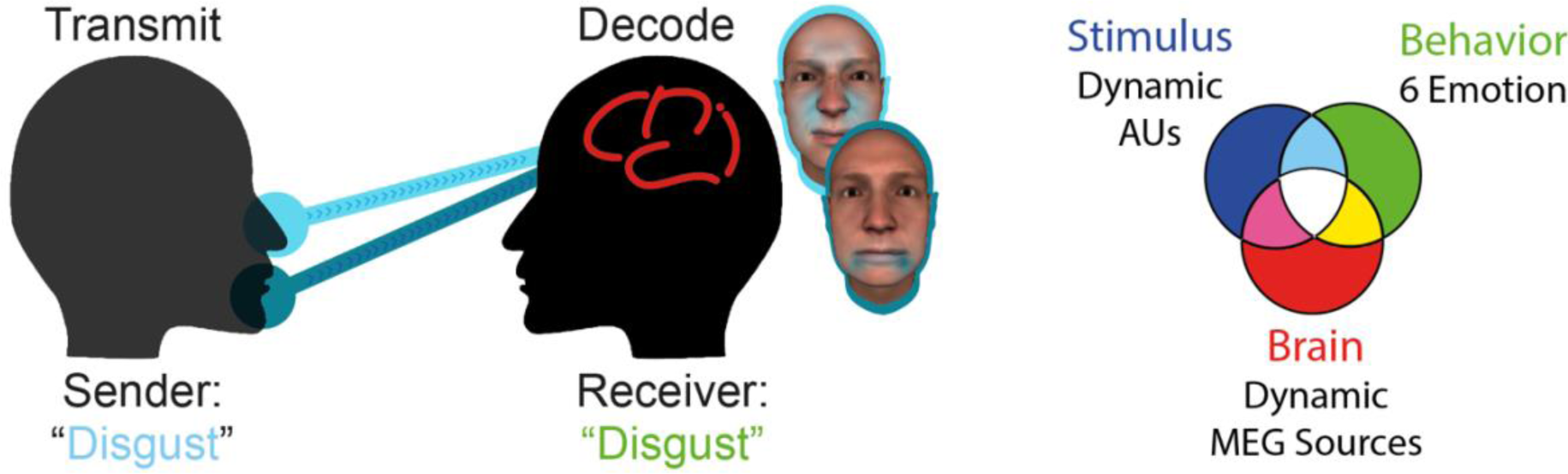
Facial signals in social interactions. The sender conveys emotion messages using different compositions of facial movements (Action Units, AUs). The receiver’s brain decodes these movements, resulting in an emotion perception–illustrated with ‘disgust.’ We uniquely study the single and double relationships (i.e. interactions) between the variables of parametrized face movement features (Action Units, AUs, blue set), participants’ behavior (categorization responses, green set) and concurrently recorded MEG brain activity (red set).

Such static and transient facial features are primarily decoded by the receiver’s brain via the ventral pathway (i.e., vision for categorization) and the dorsal pathway (i.e., vision for action), respectively^25–27^. Within the foundation of the visual hierarchy, static facial features^28^, are represented in the primary visual cortex and contralaterally projected into the ventral pathway to face-selective areas such as the occipital face area (OFA) and the fusiform face area (FFA)^29^. The dorsal pathway, known for its involvement in visuospatial processing and attention, computes the locations and general actions of objects^30,31^. Importantly, recent advances in neuroimaging have revealed in human^32,33^ and non-human primates^34,35^ what would be a third pathway that appears to process social motion^36^. This third pathway would project from occipital cortex laterally to the Middle Temporal Gyrus (MTG), the Superior Temporal Sulcus (STS), and the Superior Temporal Gyrus (STG). However, unlike the contra-lateral representations in ventral stream^29,37^, deep sources in the third pathway demonstrate ipsilateral^38,39^ engagement in the dynamic aspects of social perception, for example showing stronger neural responses to moving than static faces^38,40–42^. The separation of the third pathway from the ventral pathway is further suggested when face identification impairments in prosopagnosia are associated with reduced responses in ventral OFA and FFA, whereas preserved emotion recognition is associated with typical responses in MTG and STS^33,43–46^. The third social pathway therefore broadens the scope of visual categorization beyond the well-known ventral and dorsal pathways, prompting an understanding of specifically how it computes dynamic social information for socio-emotional perception and decision-making. That is, *how* does the social pathway, operating as a functional network, dynamically process (i.e. *represents*, *communicates* and *composes*) the dynamic facial Action Unit (AUs) that underpin socio-emotional perception and decision-making?

Our research addresses this critical explanatory gap by investigating the third pathway mechanisms engaged in perceiving and categorizing the six classic emotions—happiness, surprise, fear, disgust, anger, and sadness—from facial movements. We map out where and when these emotions are individually represented as component AUs and communicated across the brain’s pathways for emotion categorization. Furthermore, we reverse engineer how these AUs are then composed deep into the pathways to form a cohesive perception of emotion that influences behavior. Our approach is therefore designed to unravel how visual pathways process complex dynamic social signals, overcoming three major methodological challenges faced by researchers in this field:

**1. Control of Dynamic, Compositional Facial Stimuli:** Using facial expressions of emotion as an example, it is proposed that the third social pathway, which is attuned to dynamic social movements, processes them for categorization. To test this hypothesis, typical emotion neuroimaging work would use databases of static, full images and videos. However, these do not enable precise control and manipulation of the dynamic parameters of AUs in the expressing face–i.e. the specific features of facial movements. Thus, we now need to generate realistic 4D face stimuli that accurately reflect the nuanced and transient dynamics of the composition of AUs that express an emotion, separately from other static face features–e.g. those of identify-defining face shape. Generative face modelling will enable us to accurately measure how each AU and their compositions affect the brain and behavioral responses, an important step towards developing causal social neuroscience.

**2. Accounting for Individual Differences:** It is acknowledged that different individuals may use different compositions of facial AUs to perceive the same emotion ^47,48^. To preserve such individual differences, the methodological and analytical approaches must be able to discern the specific AUs that each participant utilizes to perceive each emotion, instead of assuming what these AUs are, or building an average that represents no one. Consequently, we ran a dense sampling neuroimaging design per participant ^49^, to derive a rich dataset for statistical inference of AU representations, communications and compositions. In addition, we tested each effect within-participant and estimated their Bayesian population prevalence for their replicability^50,51^.

**3. Mapping Dynamic Brain Pathways:** The final challenge lies in dynamically tracing the neural journey—identifying where, when, and how the three brain pathways process dynamic and static facial features for socio-emotion perception. Typical studies have focused on the static responses of brain regions to static stimuli ^52^. However, we now require real-time tracking of the brain’s responses to dynamic and static facial features with sufficient spatio-temporal granularity, as well as analytic approaches that can dissect the brain’s dynamic processes as they represent, communicate, and compute the facial features that support emotion categorization behavior. Such dynamic tracking of face features and their computations is a significant contribution to cognitive neuroimaging, offering an algorithmic understanding of brain function compared to traditional methods that only capture the brain’s static responses to full images or videos. Specifically, it offers a window into the compositionality of brain representations of emotions (Figure 1).

We therefore developed a methodology that addresses the above three challenges to transcend current limitations and enable mechanistic understanding of emotion processing in the brain pathways. Specifically, we used an explicit generative model of the human face that controls 3D shape and complexion plus its dynamic AUs^53^. On each trial, participants saw a random combination of AUs sampled from 42 AUs^23,53^, each controlled with temporal features and applied to a randomly generated 3D facial identity (Challenge 1). Each participant actively categorized the displayed dynamic facial expression as one of the six classic emotions–i.e. “happy,” “surprise,” “fear,” “disgust,” “anger,” “sad”—or “don’t know.” We then reverse engineered the specific composition of dynamic AUs that each individual participant used to categorize each emotion (Challenge 2). In a subsequent MEG experiment, we manipulated the perceptual availability of each individual AU of the participant’s own facial expression models while concurrently measuring how their brain dynamically responded to these AU changes during an active task. Using an information-theoretic framework^3,54^, we addressed how each participant’s brain pathways dynamically represent, communicate and compute each AU and their compositions to categorize emotions (Challenge 3). Finally, we conducted statistical analyses at the individual participant level^50,51^, with each participant’s data acting as an independent replication of the experiment. We also report the statistical Bayesian Population Prevalence (BPP)^50^, which provides the likelihood of replicating the observed effects across the population, coupled with the highest posterior density interval (HPDI) for precision in uncertainty estimation^50^ (cf. Challenge 2).

To preview our results, we found that representations of task-relevant emotions and their dynamic AU features propagate from occipital cortex, primarily to key regions involved in social perception (MTG, STS, and STG) where they are later composed to enable the disambiguation of emotions that are typically confused (e.g., disgust and anger). In contrast, representations of task-irrelevant static identity-defining shape features do not extend beyond occipital cortex to any pathway. Our findings validate the hypothesis of a functional third pathway for dynamic social signal processing, while providing a causal understanding of how the brain, at a systems level, computes complex dynamic social face information to produce emotion categorization behavior within individual participants.

## Results

### Behavioral experiment: Individual models of the six facial expressions of emotion

Here, to address Challenges 1 and 2, we conducted an experiment to model the AU compositions that each participant (*N* = 10) uses to categorize the six facial expressions of emotion (Figure 2A). A generative model of the face randomly sampled 42 AUs and their 6 temporal features–i.e. onset, acceleration, peak intensity, peak latency, deceleration, offset– each displayed on a randomly generated 3D face identity (see *Methods, Stimuli*). This produced a different animation per trial, that each participant categorized as “happy,” “surprise,” “fear,” “disgust,” “anger,” “sad,” or “other,” if they did not perceive one of the six emotions (see *Methods, Procedure*).

**Figure 2.**
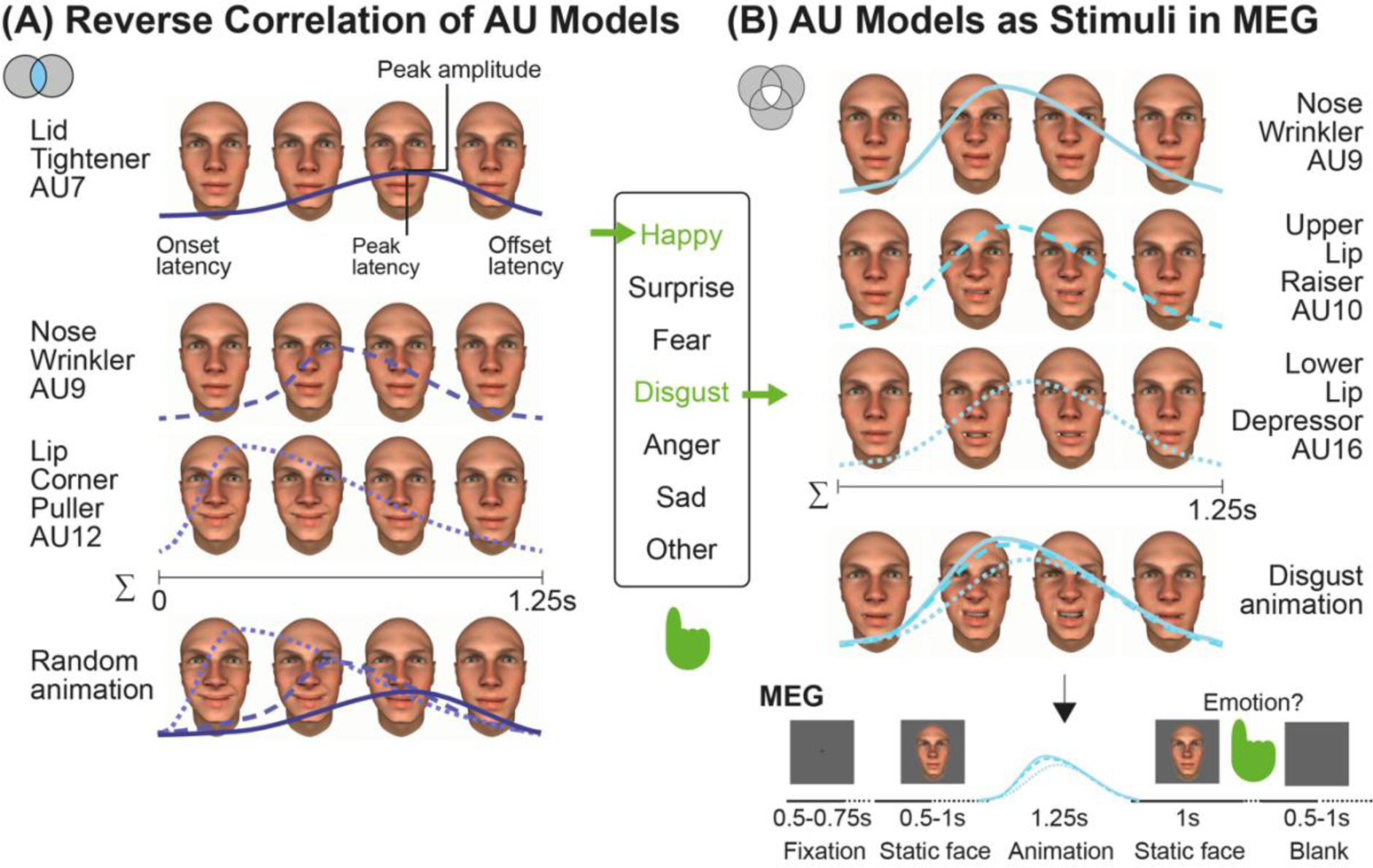
Experiment procedure. (A) Reverse correlation of AU models. On each of 2,400 trials, a generative model synthesized a facial expression animation, by randomly selecting Action Units (AUs), each animated with randomly sampled temporal parameters–onset, acceleration, peak intensity, peak latency, deceleration and offset–and displayed on a randomly generated 3D face identity, see *Methods, Stimuli*. Participants categorized these animations as one of six classic emotions (“happy”, “surprise”, “fear”, “disgust”, “anger” and “sad”) or chose “other” (i.e., 7-AFC). Following the experiment, for each emotion category we reverse correlated the AUs (two-tailed, *p*<0.05) and temporal parameters associated with behavioral responses ^53^, see *Methods, Emotion models*. For each participant, this analysis produced one dynamic model of the facial expression of each emotion. **(B) AU models as stimuli for the MEG experiment.** We used the participant’s model for each emotion to synthesize 600 participant-specific animations. In each animation, we modulated the intensity of each AU independently, setting intensity to 0 on 25% of the trials (i.e. this dynamic AU is not available in the stimulus) and varying intensity between 0 and 1 on the remaining 75% of trials. Each participant actively categorized their own dynamic models while we concurrently recorded their MEG activity over 3,600 trials, see *Methods, Participant-specific stimuli*. Each trial started with a 0.5-1s jittered static face, followed without discontinuity by a 1.25s facial expression animation displayed this time on one of 8 pre-selected 3D face identities (4 females), see *Methods, Procedure*.

Then, using each participant’s responses, we modelled the specific AUs and temporal features that caused their perception of each emotion (see *Methods, Emotion models*). Figure 3 shows the AUs of each emotion reported across all participants, displayed on the same 3D face–the cyan scale indicates the most prevalent AUs across all participants, see Supplemental Figure S1 for each participant’s emotion models.

**Figure 3.**
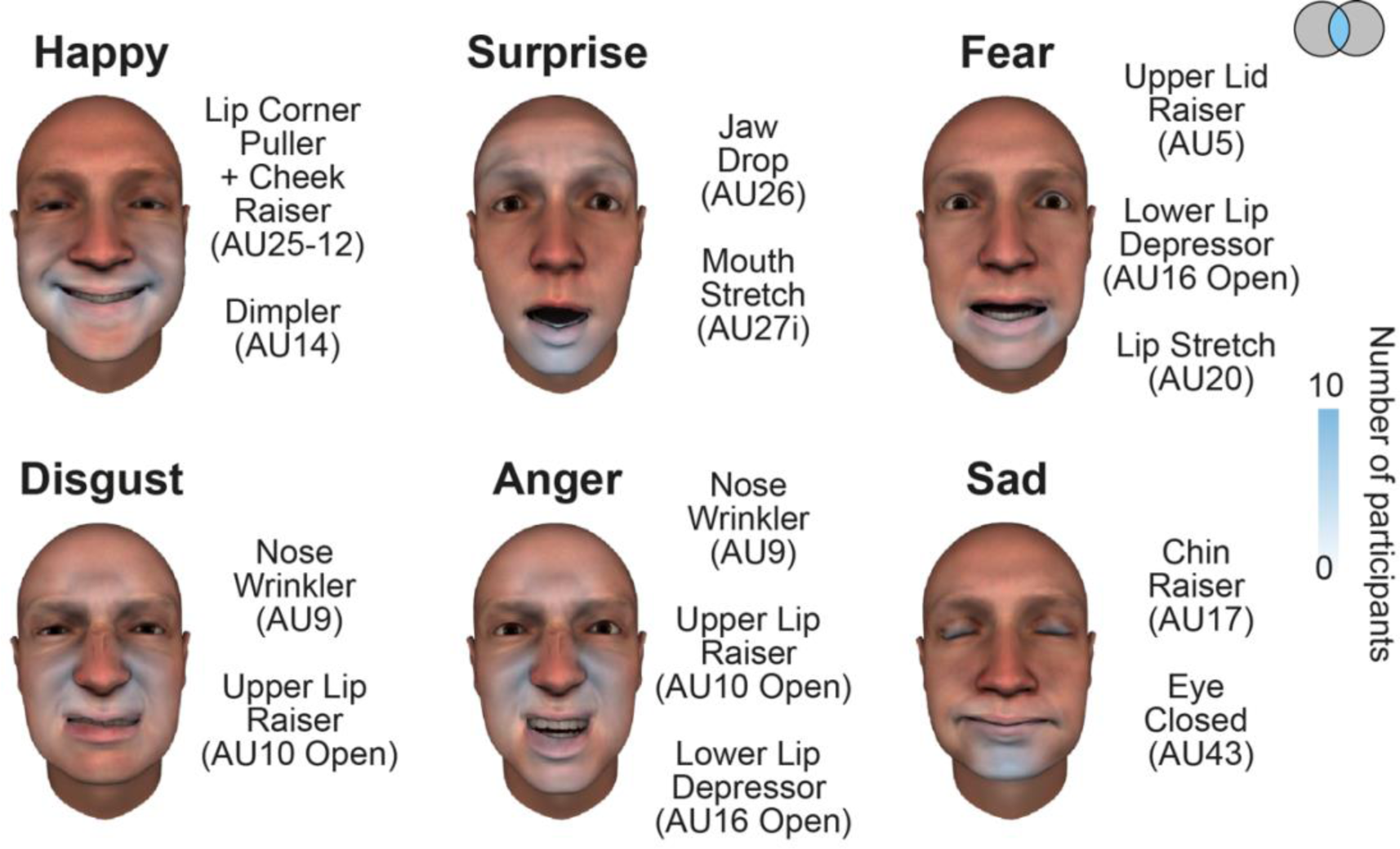
Emotion models. For each emotion category, color-coded face maps show the emotion-specific AUs across the 10 participants, see *Methods, Emotion models*. The most prevalent AUs across participants’ models are listed–i.e. *N*>=6, Bayesian Population Prevalence^50^ of the lowest boundary (i.e., N=6) = 0.6 [0.45 0.70], MAP [95% HPDI], see also Supplemental Figure S1.

### MEG experiment: Dynamic processing of the six emotions in the pathways

After modelling the specific facial movements that each participant used to categorize the six emotions, each participant returned to perform an MEG experiment in a separate session. For stimuli, we used the participant’s own facial expression models (Figure 2B and 3) because they comprise the specific AUs the participant’s brain must process to categorize each emotion. On each MEG trial, we randomly selected one facial expression model and randomly varied the intensity of each AU–i.e. set to 0 on 25% of the trials and varying between 0 and 1 on the remaining trials—thus varying the AUs’ perceptual availability. Participants actively categorized these varying animations while we concurrently recorded their brain activity measured with MEG and behavioral categorization responses.

To preview our analyses, in each participant we first reconstructed the dynamic representations of the emotion and identity categories on the MEG sources from the brain regions of the three pathways (see Table 1 for a full list of the regions included in each pathway). We then expanded this analysis to reveal the dynamic representations of each AU component from each emotion model of each participant. Finally, we tested specifically how the dynamic composition of AUs deep in the third pathway sources influences emotion perception and categorization behavior.

**Table 1.** Brain regions of the three pathways (color-coded in Figure 4).

| Pathways | Regions |
| --- | --- |
| Occipital cortex | Lingual Gyrus (LG)<br>Pericalcarine (PCAL)<br>Cuneus (CUN)<br>Lateral Occipital Cortex (LOC) |
| Ventral pathway | Inferior Temporal Gyrus (ITG) |
| Dorsal pathway | Inferior Parietal Lobe (IPL)<br>Superior Parietal Lobe (SPL) |
| Social (third) pathway | Medial Temporal Gyrus (MTG)<br>Bank of Superior Temporal Sulcus (BANKSSTS, abbreviate as STS)<br>Superior Temporal Gyrus (STG) |

### A. Dynamic representation of emotions and identities in the brain pathways

To understand how brain pathways dynamically process facial expressions of emotion, we first analyzed how each participant’s MEG activity represents the six emotion categories. To reveal these representations, we computed the Mutual Information^54^ (MI) between the varying emotion model randomly chosen on each trial (i.e. its category label) and the corresponding MEG amplitude response of the sources of the three pathways–i.e. computed as MI(emotion category; MEG_t_), every 4ms between 0ms and 500ms post animation onset, FWER-corrected over 2775 sources × 125 time points per participant, *p*<0.05. This computation produced a source-by-time matrix of MI values per participant that indicates how strongly each MEG source represents the six emotion categories at each time point, see *Methods, Analyses, Emotion and Identity representations*. In Figure 4A, the white curve summarizes these dynamic data as the per-time-point cross-participant average of maximum MI across all sources.

**Figure 4.**
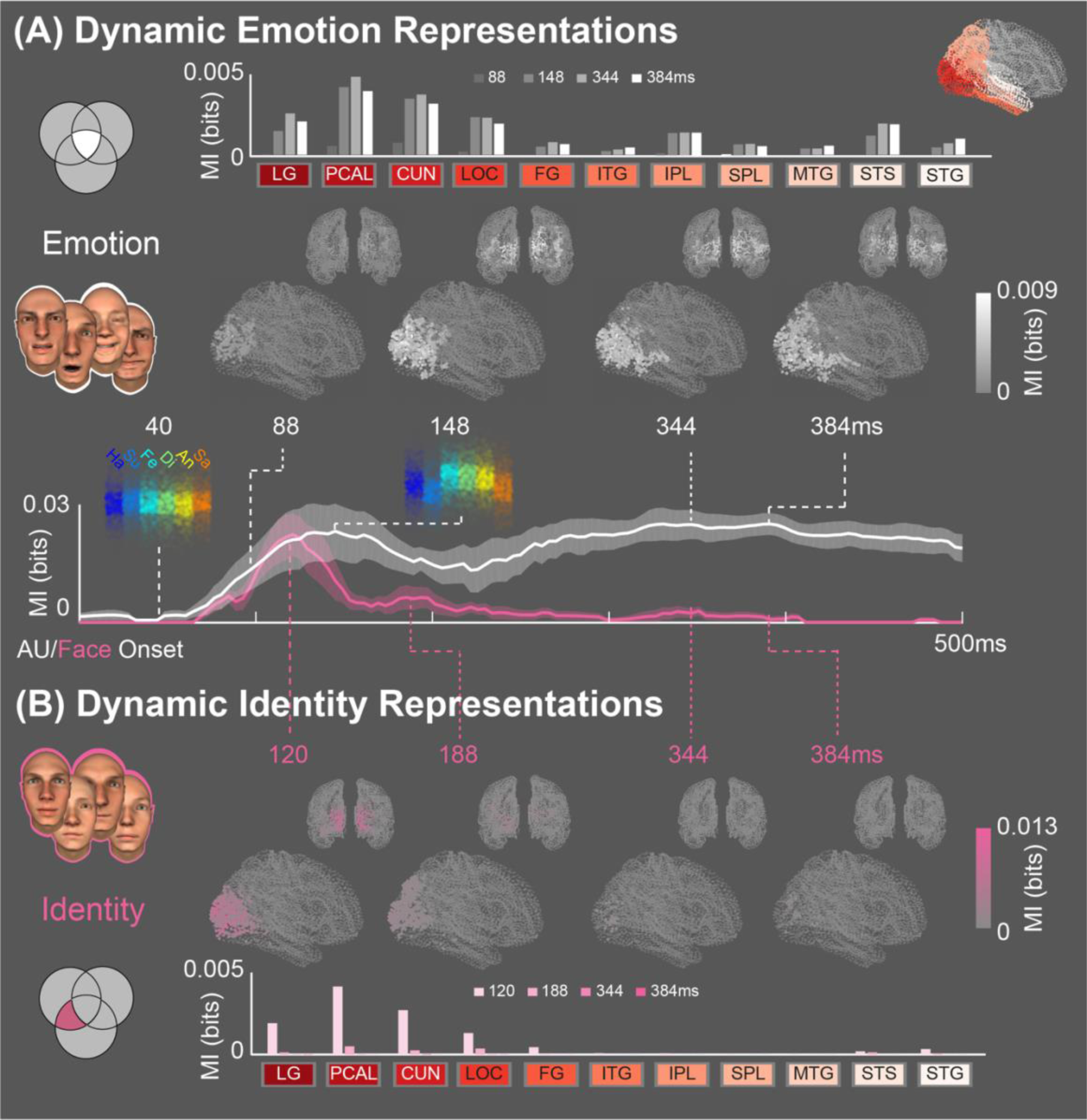
Dynamic representations of categories. (A) Emotions. We traced the dynamic representations of the emotion models, computing MI(MEG_t_; emotion category), for each source of the three pathways (see Table 1), every 4ms between 0 and 500ms post animation onset (FWER-corrected over 2,775 sources × 125 time points, *p*<0.05), see *Methods, Analyses, Emotion and identity representations*. The white curve shows the cross-participant average of the maximum MI across all these sources. To illustrate what MI represents, we color-coded by emotion model the single-trial MEG amplitude responses of an example source, at 40ms (when MI is low because MEG responses do not distinguish between the emotions) and 148ms (when MI is high because MEG responses distinguish between the emotions).The white brains show the dynamic representation of emotions at 88ms, 148ms, 344ms and 384ms post animation onset. Bar plots show the mean representation across sources in each region of the pathways (color-coded in the reference brain) and at each selected time point (early to late, from left to right). **(B) Identity.** As a control, we repeated this representation analysis using task-irrelevant face identity. We computed MI(MEG_t_; face identity label), every 4ms between 0 and 500ms from face onset (FWER-corrected over 2,775 sources × 125 time points, *p*<0.05). The magenta curve shows the cross-participant average of maximum MI across sources. The magenta brains show identity representation at 120ms, 188ms, 344ms and 384ms. Bar plots show the mean representation across sources in the regions of the pathways and at each selected time point (early to late, from left to right). See also Supplemental Figures S1 and S2.

At this juncture, we pause to emphasize that the MI curve does *not* show an average response of the brain to the six emotion categories. Instead, it is a direct measure of how strongly the variations of MEG source response represent the six emotion categories–i.e. quantifying how well the MEG responses discriminate the emotion stimuli (e.g. see color-coded inset in Figure 4A, at 40 ms when sources do not yet discriminate the emotions, and at 148 ms when they do). White brains and bar plots above localize the dynamic flow of these emotion representations at four time points. They reveal early representations in occipital cortex (at 88 ms), from which the flow primarily progresses laterally, along the social pathway (from 148 ms) to MTG, BANKSSTS (henceforth, abbreviated as STS) and STG (344-384 ms). Emotion representations also flow in the dorsal pathway to IPL and SPL, but these do not ventrally extend to ITG typically associated with face processing^38^. We replicated the significant emotion representation in each individual participant^50^–Bayesian Population Prevalence = 1.0 [0.75 1.0], MAP [95% HPDI], and the significant difference between ventral and social pathways in 9/10 participants, BPP = 0.9 [0.61 0.99], MAP [95% HPDI], see Supplemental Figure S1 for pathway representations of each individual emotion model. We also replicated this representational pathway per emotion, see Supplemental Figure S2.

To compare with the dynamic representations of facial expressions, we similarly computed the representation of the 8 task-irrelevant face identities conveyed by the static 3D face shape features prior to the onset of participants’ facial expressions–i.e. computed as MI(face identity; MEG_t_), FWER-corrected over 2775 sources × 125 time points, *p*<0.05. The magenta curve in Figure 4B also shows an early representation of static face identity in occipital cortex (at 120 ms), but this time followed by its quick reduction, with no further propagation into higher pathways regions^55,56^ (see magenta cortical and bar plots in Figure 4B). We replicated this result in each individual participant–BPP = 1.0 [0.75 1.0], see Supplemental Figure S1.

### B. Dynamic AU representations in the social pathway

Having shown that the brain dynamically represents emotions along the third social pathway while reducing identity representations in occipital cortex, we now focus on the AUs that comprise these brain representations. Remember that our MEG experiment randomly varied the intensities of each AU of each participant’s facial expression models. Thus, for each modelled AU, we can compute across trials how MEG source amplitude responses dynamically represent AU intensity variations–i.e. computed as MI(MEG_t_; AU intensity), every 4ms between 0ms and 800ms, FWER-corrected over 2,775 sources × 200 time points, *p*<0.05, see *Methods, Analyses, AU representation*. The color-coded MEG amplitudes insets in Figure 5A illustrate the representation stages of one AU, where the MEG of one STS source does not represent the intensity variations of the nose wrinkler AU at 40 ms, but does so later, at 370 ms. Figure 5A summarizes these AU representation results across participants. The white curve shows the average time course of the per-participant AU representations–i.e. it is computed for each participant and AU as the per-time-point maximum MI across sources, then averaged across all AUs and participants. Together, these results confirm that the sources of the regions forming the third social pathway do indeed dynamically represent the AUs that support emotion categorization behavior, from occipital cortex to MTG, STS and STG. We replicated this result in each individual participant^50^–BPP = 1.0 [0.75 1.0], MAP [95% HPDI], and reported them in Supplemental Figure S3.

**Figure 5.**
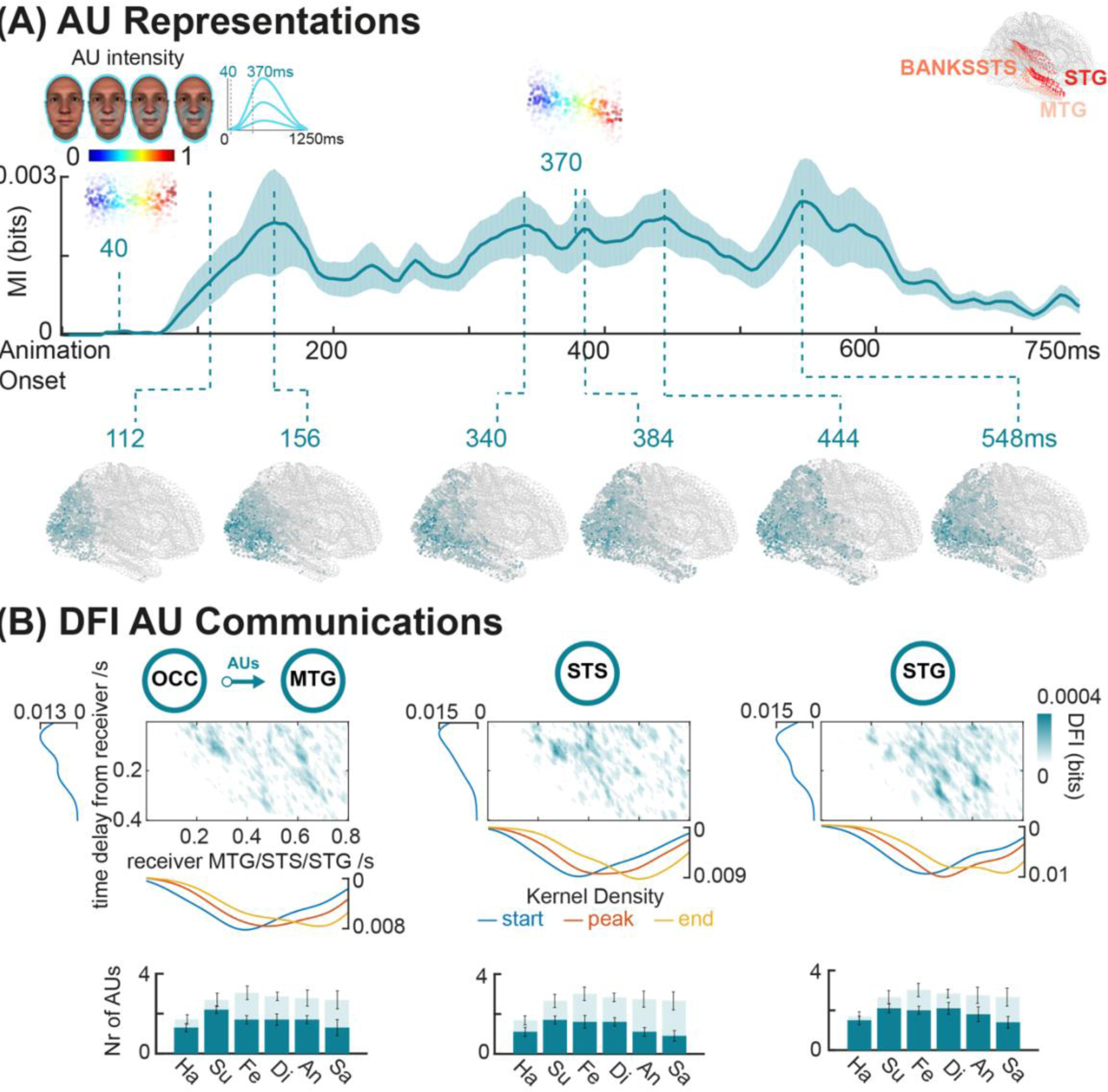
Dynamics of AU representations and communications. (A) Representations. For each participant and emotion model we computed the representational dynamics of each AU as MI(MEG_t_; AU intensity, FWER-corrected over 2,775 sources × 200 time points, *p*<0.05, see *Methods, Analyses, AU representation*. Across all participants and AUs, the curve shows the mean of maximum MI across ventral, dorsal and social pathways sources (Table 1), localized at six different time points. The color-coded inset illustrates single-trial MEG amplitudes of an STS source responding to the variations of intensity of nose wrinkler AU in Participant 5, when MI is low (at 40 ms) vs. high (at 370 ms). **(B) Communications.** For each participant, emotion model and AU, we selected a sender (in occipital cortex) and receiver pair of sources (independently in MTG, STS and STG) with maximum significant MI. For each pair, we computed DFI communications between sender and receiver sources every 4 ms between 0 and 800ms post animation in the time course of the receiver (x-axis), every 4ms delay between 0 and 400ms from the sender (y-axis), see *Methods, Analyses, AU communication*. <u>Matrix</u> <u>plot</u> reports DFI communications separately to MTG, STS and STG, as the cross-participant and emotion models mean of significant DFI–FWER across 200 receiver time points × 100 delays, *p*<0.05, across all participants and AUs. <u>Curves</u> show the kernel density estimation of the start of DFI communications (as first receiving time point, blue), peak (orange) and end (last receiving time point, yellow). <u>Bar plots</u> show the number of significantly represented AUs per emotion in occipital cortex (light) which are significantly DFI-communicated to MTG, STS and STG receivers (dark). See Supplemental Figure S3 for individual participants data.

### C. AU communications in the social pathway

The dynamic flow of AU representations suggests a pathway functionally extending from occipital cortex to MTG/STS/STG that specifically represents and communicates AUs for social categorization. Formally establishing this functional property requires quantifying specifically the communications of AUs across sources from all other possible communications. We did so using Directed Feature Information (DFI)^57,58^, which can quantify directed, time-lagged, source-to-source communications of AU features, see *Methods, Analyses, AU communication*.

In each participant, we therefore identified the potential pairs of sending (occipital) and receiving (MTG/STS/STG) sources that maximally represent each AU. For each source pair, we then computed the AU communication as DFI_AU_(sender_OCC_ -> receiver_MTG/STS/STG_) ^57,58^, using communication delays between 0 and 400 ms, FWER-corrected over 200 receiver time points × 100 delays, *p*<0.05. Figure 5B shows the resulting communication matrix, as the cross-participant average of all DFI_AU_(sender_OCC_ -> receiver_MTG/STS/STG_). At least one AU was significantly communicated from OCC to MTG/STS/STG in each individual participant, Bayesian Population Prevalence = 1 [0.75 1.0], MAP [95% HPDI], reported in Supplemental Figure S3.

Figure 5B indicates that MTG/STS/STG sources receive AUs sent from occipital sources between 200 to 700ms post animation onset. Kernel density estimates indicate the distributions of start, peak and ending points of these communications within the period. For each emotion, the histograms below show the number of AUs that occipital cortex represents and then communicates to MTG/STS/STG.

### D. Source-level representations of AUs discriminate categorization behavior

In an active emotion categorization task, we showed that the social pathway communicates the individual AUs of facial expression models from occipital cortex to MTG/STS/STG sources. Now, we turn to understand how the sources that receive these AUs via the social pathway specifically compute (i.e. compose) them for categorization behavior. We focus on the well-established confusions between disgust and anger—two similar facial expressions—which can only be discriminated when different AUs are composed ^24^.

Figure 6A shows that Nose Wrinkler AU, which is shared between anger and disgust, leads to either “disgust” or “anger” categorization behaviors (Figure 6A, Behavior, light green). This is because Nose Wrinkler, on its own, cannot discriminate “disgust” and “anger” (Behavior, dark green). However, adding to Nose Wrinkler either Lip Stretcher, Lip Corner Puller, or Lower Lip Depressor AUs leads to successful discriminations of these emotions (Figure 6A, Behavior, dark green).

**Figure 6.**
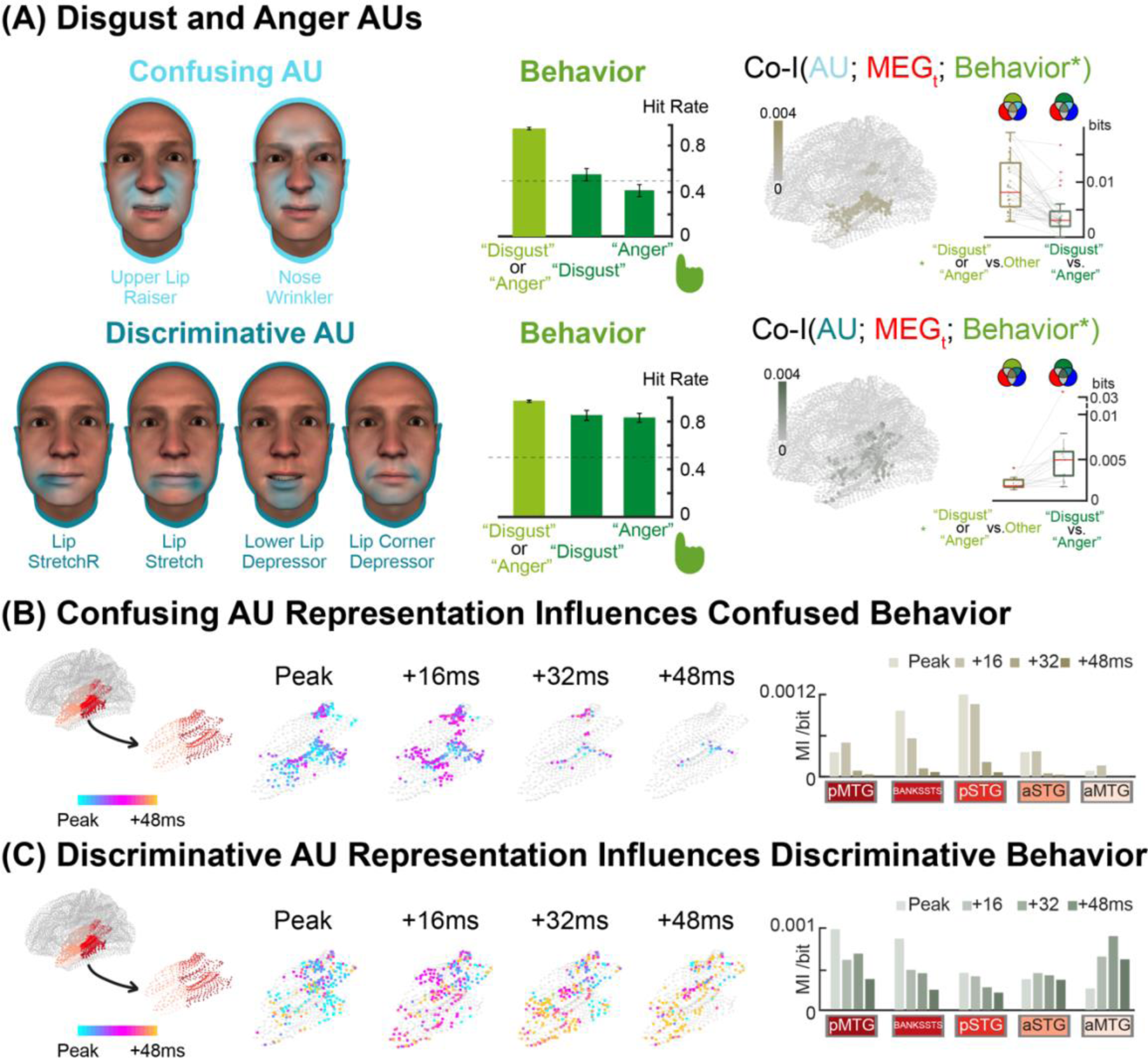
AU compositions disambiguate behavior. (A) Disgust and Anger AUs. Disgust and anger are behaviorally confused because they share Nose Wrinkler and Upper Lip Raiser AUs. <u>Behavior.</u> Bar plots (±standard error) show these behavioral confusions when confusing AUs comprise the stimuli, leading participants to respond either disgust or ‘anger with highly probability (light green) but confusing disgust and anger animations (dark green). Adding the discriminative AUs Lip Stretcher, Lip Corner Puller, or Lower Lip Depressor to the ambiguous Nose Wrinkler and Upper Lip Raiser (dark green) disambiguates disgust and anger categorizations. <u>Co-I.</u> For each confusing and discriminative AU, we computed a new representational interaction with Co-Information, see *Methods, Analyses, AU modulates behavior*. Specifically, we computed the overlap between the intensity variations of confusing AUs across trials (blue), corresponding MTG/STS/STG source responses (red) and behavioral categorizations (green), separately for when behavior is confused (disgust or anger vs. other emotion, light green) or correctly discriminated (disgust vs. anger, dark green). Small brains localize the sources with significant representational Co-I at peak time (cross-participant mean), when behavior is confused vs. discriminated. At this representational peak, box-plots show source Co-I distributions across participants when categorization behavior is confused vs. discriminated. **(B) Confusing AU representations confuse subsequent categorization behavior.** Color-coded MTG, STS and STG sources show peak time Co-I representation of confusing AUs when behavior is confused. Co-I then falls off from in 16 ms time steps in a representational dynamics primarily located in posterior MTG, STG and BANKSSTS (i.e., pSTS). Bar plots show mean Co-I across sources in each region and each time point. **(C) Confusing plus discriminative AUs representations discriminate subsequent categorization behavior.** Same analysis but when Co-I comprise representation of discriminative AU and discriminated behavior. The representational dynamics this time shows transitions from pMTG, pSTS to aMTG 48ms following the peak.

In Supplemental Figure S4, we first establish that the MEG amplitudes of single MTG/STS/STG sources compose confusing and discriminative AUs–i.e. which are synergistically represented on the same sources, see *Methods, Analyses, AU compositions*. Amongst MTG/STS/STG sources, we localize those leading to confusions vs. discriminations of “disgust” and “anger.” To do so, we separate the trials showing “disgust” and “anger” animations into those comprising the confusing Nose Wrinkler AU (i.e., leading to “disgust” and “anger” confusions) from those comprising <u>both</u> the confusing and discriminative AUs (i.e., whose composition discriminates “disgust” and “anger”). We use these confused vs. discriminated trials to demonstrate how AU representations at source-level disambiguate emotion categorizations.

To do so, we interlock the three key variables of our MEG design: AU intensity in stimulus animation, STG/STS/MTG source amplitude response and participant’s emotion categorization behavior, using the distinction between confused vs. discriminated trials. Specifically, we compute a new measure of representational overlap–i.e. of AU intensity and categorization behavior into the amplitude of individual STG/STS/MTG sources. This double interaction is quantified with Co-Information (Co-I), as the information shared between AU intensity and categorization behavior into MEG amplitude, see *Methods, Analyses, AU modulates behavior*. Thus, with Co-I we identify which sources significantly represent confusing AUs when “disgust” and “anger” are subsequently confused (Figure 6A, Co-I panel); and which specific sources represent discriminative AUs when behavior subsequently discriminates the two emotions (Figure 6A, Co-I panel).

Each Co-I panel in Figure 6A localizes with a small brain the AU-representing sources leading to behavioral confusions (top panel) and those leading to behavioral discriminations (bottom panel). A group-level two (confusing vs. discriminative AUs represented) by two (representational interactions with confused vs. discriminated categorizations) ANOVA between the bar plots of the top and bottom panels reveals a significant interaction, *F*_(1,92)_ = 17.46, *p* < 0.0001. This implies that these sources represent different AUs for different categorization behaviors.

The inset in Figures 6B and C detail the dynamics of these representational interactions. Figure 6B shows that significant representational interactions of confusing AUs leading to confused “anger” and “disgust” behaviors are primarily confined to the posterior sources of MTG, STG and BANKSSTS in pSTS–FWER-corrected over 661 sources, *p*<0.05, and replicated for at least one confusing AU in 9/10 participants, BPP = 0.9 [0.61 0.99], MAP [95% HPDI]. In contrast (Figure 6C), when the animations show both a discriminative AU (e.g. lower lip depressor) and a confusing AU (e.g. Nose Wrinkler), that participants correctly discriminate as “disgust” and “anger,” representational interactions dynamically evolve from pMTG, pSTS to aMTG over 48ms post representation peak–FWER-corrected, *p*<0.05, and replicated for at least one discriminative AU in 9/10 participants, BPP = 0.9 [0.61 0.99], MAP [95% HPDI].

Together, these analyses show that deep social pathway sources differentially represent (in space x time) AUs and their combinations to disambiguate subsequent behavioral categorizations of facial expressions of emotions.

## Discussion

We aimed to answer the critical questions of where, when and how brain pathways process dynamic social signals to categorize them, using the pervasive facial expressions of the six classic emotions. We proceeded in two steps. First, we coupled a generative model of facial movements (AUs) with each individual participant’s categorization behavior to reverse engineer a dynamically parametrized model of the specific AU features that drive emotion categorization behavior in each participant. Next, we measured each participant’s MEG responses to their own six facial expression models where the intensity of their AU features was randomly varied across trials. Information theoretic source-level analyses revealed a functional, third social pathway–beyond the well-known ventral and dorsal pathways–that dynamically processes facial movements for categorization behavior. Specifically, its sources represent the task-relevant AUs features of the participant’s emotion models, communicates these AUs between occipital cortex and MTG, STS and STG, where deep sources compute–i.e. compose–AU features to disambiguate categorization behavior. In contrast, task-irrelevant 3D identity features are initially represented but then swiftly reduced in occipital cortex. Building on lesion studies that suggested a functional separation between the ventral and social pathways, our study provides a computational understanding of <u>how</u> individual participants’ brain, at a systems level, dynamically computes complex dynamic social signals to produce emotion categorization behavior.

Beyond facial expressions, the third social pathway also responds selectively to multiple other social signals such as body ^59–62^ and speech features ^63–65^, which are integral to human social interactions. Our new representational interactions analyses revealed that STS sources compose different AUs received at different time points to disambiguate categorization behavior. This raises the important question of the multi-modal nature of social perception and whether the disambiguation that we demonstrated straightforwardly extends to computations of social signals across different signaling modalities–e.g. where the higher-pitched feature of a happy voice could be composed together the AU features of the facial expression to facilitate categorization of a ‘happy’ face^66^. Our approach suggests a new way to study such multimodal interactions. In a first step, we would use generative models of the voice (or the body) and model the representations of their features for a variety of social categorization tasks–e.g. emotion categories, but also valence and arousal ^14^, trustworthiness ^67^, social dominance ^68,69^, attractiveness ^12^, emotion intensity ^70^ and so forth. Such studies should be sensitive to the culture of each individual participant ^12,48^. Next, we would bring back the participants and independently vary their individual models in each modality (e.g. visual for face and/or body features and auditory for voice features) to study the scope of feature representation, communication and computations across modalities in the corresponding pathways.

It is often assumed that the brain selectively processes sensory features for visual categorization. However, controlling the stimulus features (instead of using large image/video databases) is required to understand the selective features that brain pathways compute for behavior ^71,72^. As such control is rare, the computational processes underlying selective feature processing remain unclear. Here, as in^56^, such a feature control revealed a rapid occipital representation then reduction of the task-irrelevant features–but here, with 3D face identity features, all reduced within 170 ms post stimulus. In contrast to ^56^, where we showed that communications of task-relevant features to rFG sources in the ventral pathway, we showed that communications of dynamic AU features selectively to STG/STS/STG sources in the third pathway. This finding suggests new research avenues. The first one concerns face familiarity. If the 3D face identity features represented a familiar face^73^, they could be passed into the rFG, instead of reduced in occipital cortex, even though the active emotion categorization task would require processing the dynamic AU features. Similarly, switching the active task to categorizing the identity of these same dynamic faces could lead to the reduction of their AU features in occipital cortex. Further research should seek to understand the neural mechanisms of task-guided face processing, how the brain flexibly reprograms to use different features from the same faces depending on task demands, and how such flexibility influences conscious stimulus perceptions–e.g. a familiar face in the context of an active emotion categorization task could be the visual analogue of hearing one’s own name in the auditory cocktail party effect ^74,75^.

Though we focused on the three visual pathways, we know that higher hierarchical brain regions responsible for executively control could influence feature representation, communication and composition in a top-down manner ^58,76^. Previous research suggested involvement of anterior cingulate cortex ^77,78^, medial prefrontal cortex ^79^ and amygdala ^80^ to orchestrate attention control (including to stimulus features ^55,58^) and modulate the neural activity of primary visual cortex. Furthermore, we also noticed an earlier dorsal pathway engagement with AU representations in IPL and SPL ^81^, which could contribute to broader context facilitating the engagement of the social pathway with dynamic AUs. Future research should specifically investigate the influence of other brain regions, such as prefrontal cortex, anterior cingulate cortex, or insula and amygdala, on AU representation, communication and computation in the social pathway.

Our innovative methodology paves the way for a computational approach to social neuroscience that rigorously explores how humans perceive, categorize, and interact with the complex social world. Central to our approach is the analysis-by-synthesis framework ^82–85^, which posits that we can only understand the visual world through the lens of what we can recreate or model generatively. This framework deconstructs the visual world’s complexity into simpler component features, resonating with neuroscientific theories that trace the processing of basic Gabor edges in V1 to surfaces in higher cortical areas. Therefore, the central problem of AI and computational cognitive neuroimaging of aligning the human brain to its models, implies that both should process the same features with the same algorithmic computations to yield similar behaviors ^71,72^. By adopting a componential modelling approach, we facilitate a shift from treating the brain and our models as opaque (“black boxes”) to more transparent “glass boxes.” This transparency enables us to directly link specific behaviours to distinct computational processes in the visual system, enhancing our understanding of the underlying mechanisms of social cognition.

To conclude, using a generative model to manipulate individual AUs, we showed the third social pathway functionally represents, communicates and composes AUs for emotion categorization, providing a mechanistic understanding of socio-emotional perception in the brain. This approach is broadly generalizable to develop a new computational social neuroscience to investigate how we perceive, interpret, and navigate the multifaceted social world.

## Methods Participants

Ten participants (6 female, Western Caucasian, 18-35 years, mean=25.1, SD=3.4) took part in the experiment and provided informed consent. All had normal or corrected-to-normal vision and reported no history of disorders related to processing facial signals (e.g., autism spectrum disorders), faces (e.g., prosopagnosia), learning difficulties, synaesthesia, or other psychiatric/neurological disorders. The University of Glasgow College of Science and Engineering Ethics Committee approved the experiment (Application Number: 200220190). Each participant first completed the behavioral experiment before being transferred to the subsequent MEG experiment, as explained below.

## Behavioral experiment

### Stimuli

We used a generative model of human facial movements^53^, which is comprised of a library of 42 individual facial action units (AUs)—i.e., the basic elements of human facial movements as detailed by the taxonomic Facial Action Coding System (FACS)^23^. Each AU in the generative model was derived from real humans, trained to accurately produce each individual AU on their face, captured using a stereoscopic system and their rendering verified by the trained AU producer^53^. Therefore, the generative model produces valid representations of real human facial movements and comprises no physiologically impossible facial movements. To generate facial expression stimuli on each experimental trial, the generative model randomly selected a combination of AUs from a set of 42 (minimum = 1 AU, maximum = 6 AUs, median = 4 AUs selected across trials) and assigned a random movement to each AU from six temporal parameters whose values were randomly chosen—onset latency, acceleration, peak intensity, peak latency, deceleration, offset latency. These six temporal parameters enabled each AU to peak once during the 1.25s stimulus time course, while other parameters such as acceleration and intensity could vary across the experiment, thus enabling the exploration of dynamic properties while retaining experimental feasibility^53^.

Using this generative model, we generated 2,400 random facial expression animations and displayed each on a different and randomly generated face identity using a 3D face identity generator ^73^ (Western Caucasian, 1,200 females, 1,200 males aged 20–40 years). All facial animations were presented in the center of the participant’s visual field, on the grey background (RGB: 100, 100, 100) of a 19-inch flat panel Dell monitor (Round Rock, Texas 78682, refresh rate of 60 Hz and resolution of 1024×1280). Participants used a chin rest to maintain a constant viewing distance of 70cm, with stimuli subtending 8° (vertical)×6° (horizontal) of visual angle.

### Procedure

On each trial, participants viewed the randomly generated facial animation that they categorized according to one of the six classic emotions by pressing dedicated response keys—i.e., ‘‘happy,’’ ‘‘surprise,’’ ‘‘fear,’’ ‘‘disgust,’’ ‘‘anger’’ or ‘‘sad’’^86^. Participants could only correctly categorize an animation when it matched one of the six emotions—i.e., corresponded with their prior knowledge of one of the 6 facial expressions of emotion. If the facial movement did not represent any of the emotions, participant selected ‘‘don’t know.’’ We counterbalanced response keys across participants.

Each trial started with an animation played once for 1.25 s followed by a blank screen. We instructed participants to respond as quickly and as accurately as possible by pressing the appropriate key, which they could only do after the animation had finished (and within 5 s post animation). The next trial started following response. We divided the randomized trials into 24 blocks of 100 trials. After eight consecutive blocks of 100 trials, participants took a break of at least 1 hour.

### Emotion models

For each participant, we used an established model-fitting procedure^53^ to compute a model for each of the six emotions. Briefly, we first identified the AUs associated with each emotion category. That is, for each emotion, we Pearson correlated two binary vectors whose cells correspond to the experimental trials. The cells of the first vector represent the presence vs. absence of a given AU on each trial; Those of the second vector represent the corresponding categorization responses–i.e. hit vs. not-hit categorization of this emotion. For each emotion category this analysis delivered the 42-dimensional vector of the AUs significantly correlated (two-tailed, *p*<0.05) with categorizing this emotion. Then, for each significant AU in the vector, we further modelled its dynamics by linearly regressing the hit vs. not-hit response vector with each one of the six temporal parameters. These analyses produced a model–i.e. AUs and their dynamic parameters–describing the dynamic facial expression that this participant perceives at that emotion (summarized in Figure 3).

## MEG experiment

Each participant returned to perform an MEG experiment in a separate session.

### Participant-specific MEG stimuli

In the MEG experiment, we used each participant’s emotion models to synthesize 600 new animations that mapped onto 8 generated Western Caucasian 3D face identities (4 males). These animations modulated the perceptual availability of each AU by randomly modulating its intensity parameter while the dynamic parameters of other AUs remained unchanged. Specifically, on 25% of the trials, intensity was set to 0; on the remaining 75% intensity randomly varied between 0 and 1. We produced a total of 3,600 animations per participant (600 animations x 6 per-participant emotion models). In the MEG scanner, participants maintained a constant viewing distance of 115cm, with stimuli subtending 8° (vertical)×6° (horizontal) of visual angle.

### Procedure

On each trial, participants viewed a randomly selected animation that they categorized using the same response options as in the behavioral task–the six emotion categories plus “don’t know.” Each trial started with a 0.5s to 1s static face with jitter. Then, a 1.25s animation played once, immediately followed by a 1.25s to 1.8s inter-trial interval (ITI) with jitter. A fixation cross was maintained in the centre of the screen throughout the experiment. Each participant completed a total of 72 blocks of 50 trials, spread across 4-5 sessions run over 4-5 days, for a total of 3,600 trials per participant.

## MEG Data Acquisition and Pre-processing

We measured each participant’s MEG activity with a 306-channel Elekta Neuromag MEG scanner (MEGIN) at a 1,000Hz sampling rate. We performed the analyses according to recommended guidelines using the MNE-python software^87,88^ and in-house Python/MATLAB code.

We rejected noisy channels with Maxwell filtering and visual inspection, and blocks of trials with a head movement > 0.6cm (tracked by cHPI measurement). For each remaining block, we applied signal-space separation (SSS)^89,90^ to the raw data to reduce environmental noise and compensate for head movement. We band-pass filtered the data between 1-150Hz (Hamming FIR filter), notch-filtered them at 50, 100 and 150Hz and rejected muscle artifacts with automatic detection. We separately epoched the output data into [-200ms to 500ms] trial windows around static face onset and [-200ms to 800ms] trial windows around face movement onset and rejected jump artifacts with automatic detection. We concatenated the epoched data of all blocks per session, decomposed the output dataset with ICA, identified and removed the independent components corresponding to artifacts (eye movements, heartbeat—i.e., 2-5 components/participant).

We resampled the output data at 250 Hz, low-pass filtered them at 25Hz (5^th^ Hamming FIR filter) and performed the minimum-norm estimate (MNE) analysis with an empty-room recording. We reconstructed the time series of MEG sources on a 5mm grid of boundary element model (BEM) surface (computed with Freesurfer and MNE software per participant). We applied this reconstruction to each session of trials. These computations produced for each participant a matrix of single-trial MEG response time series–of dimensions 8,196 MEG sources x 250Hz sampling rate.

## Analyses

### Emotion and identity representations

To evaluate the dynamic representation of the emotion and identity categories, we proceeded as follows. For each participant, we computed Mutual Information (MI) as MI(6 emotion labels; MEG_t_) and MI(8 identities; MEG_t_) every 4ms from 0 to 500ms following static face onset (for identity), and following animation onset (for emotions), on each of 2,775 sources of the ventral, dorsal and third pathway–i.e. lingual gyrus, pericalcarine, cuneus, lateral occipital cortex, fusiform gyrus, inferior temporal gyrus, middle temporal gyrus, superior temporal gyrus, inferior parietal lobe and superior parietal lobe, see Table 1.

We computed MI with the Gaussian Copula Mutual Information (GCMI) estimator^54^. For statistical testing, we repeated the above MI computations 200 times, while shuffling either the identity or emotion labels at each iteration and extracted the maximum MI value across the MI source x time matrix of each shuffled repetition (FWER, *p*<0.05, one-tailed). We used these 200 maxima to establish the statistical threshold as the 95th percentile of the distribution.

We averaged across participants the maximum MI across sources at each time point, separately for emotion and identity representations. Figure 4 plots these means representation curves and locate them in the brain at four different time points post stimulus.

### AU representation

To investigate dynamic AU representations in the brain, we used each participant’s emotion models and computed, for each AU, the cross-trial MI(AU intensity; MEG_t_) on all 2,775 sources, every 4ms between 0ms and 800ms (FWER-corrected, *p*<0.05). To establish statistical significance, for each AU we repeated 200 times the MI computations with shuffled intensities, using the 95th percentile of the distribution of 200 maxima as threshold–each maximum taken across the source x time MI matrix of each shuffled repetition, FWER, *p*<0.05, one-tailed. We then took the maximum AU representations across sources at each time point, and averaged the output time courses across AUs and participants. Figure 5A shows these results.

### AU communication

To reconstruct the communications of each AU along the social pathway, we computed Directed Feature Information (DFI, where F is the AU intensity) between occipital and MTG/STS/STG as follows.

*<u>Step 1: Source selection</u>*. For each AU, in each emotion model, we selected the sender (in occipital cortex) and the receiver (independently in MTG, STS and STG) with maximum significant MI.

*<u>Step 2: Directed information.</u>* Directed Information (DI, i.e., event-related Transfer Entropy) ^57,58^ quantifies all the information communicated from sending to receiving sources, removing information sent from the receiver itself. For the receiver at time *x*, with a communication delay *y* from the sender, DI is computed as the conditional mutual information between *RA_x_* and *SA_x-y_* conditioned on *RA_x-y_* as follows:

DI= CMI <*RA_x_*; *SA_x-y_*| *RA_x-y_*>

Thus, we computed DI between the sender and receiver sources for each receiver time point between 0 and 800 ms post animation onset and for each communication delay between 0 and 400ms. This produced the receiver-time x transfer-delay DI matrix.

*<u>Step 3 DI conditioned on feature (DI|F)</u>*: DI|F removes from DI the information communicated about the AU intensity itself. We computed DI|F for each receiving time x communication delay.

*<u>Step 4 DFI:</u>* The difference between DI and DI|F therefore isolates the information communicated between sending and receiving sources about the AU. We computed DFI as follows:

DFI = DI– DI|F

for each receiving-time x communication-delay cell of the matrix.

*Step 5 Statistical significance*: We repeated 200 times DFI computations with shuffled AU intensities, using as the statistical threshold the 95th percentile of the distribution of 200 maxima (each taken across the DFI matrix of each shuffled repetition).

We replicated these steps for each AU in each participant’s model and averaged the output DFI matrices across AUs as shown in Figure 5B. We plotted the kernel density estimation of the communication start (first received point), peak and end (last received time point) across all AUs in all participants. Additionally, for each of six emotions, we plotted the AUs significantly represented (in occipital cortex) across all participants and those significantly communicated AUs (to STG/MTG/STS).

### AU composition

To understand when, where and how MTG/STS/STG sources compose AUs following their communications, we computed the synergistic representation of pairs of AUs into MEG source activity. Specifically, for each combination of a confusing AU (i.e. Nose Wrinkler and Upper Lip Raiser) and a discriminative AU (i.e. Lip Stretch, Lip Corner Puller and Lower Lip Depressor) in the participant’s models, we computed the Co-information between confusing AU intensity, discriminative AU intensity and MEG_t_ activity as follows:

Syn = -(MI(confusing AU intensity; MEG_t_) + MI(discriminative AU intensity; MEG_t_) -MI(confusing AU intensity, discriminative AU intensity; MEG_t_))

Across all participants and AU pairs, we averaged these synergistic representations and aligned them at the peak representation time of the second AU. Supplemental Figure S4 shows the group mean. To illustrate these synergistic representations, for each AU pair, we averaged the MEG activity of the source representing both AUs from -80ms to 80ms around the representation peak (MI) of the second AU. We report three group-level conditions in Supplemental Figure S4: trials when the confusing AU is absent from the animation (intensity value == 0), when this AU is present and when it is present together with the discriminative AU.

### AU modulates behavior

To understand when, where and how source representations of AUs modulate subsequent decision behavior, for each participant we computed the representational overlap between AU intensity, behavioral decisions and MEG activity of MTG/STS/STG sources as follows:

On trials showing disgust and anger stimuli that present the confusing AUs, we computed for each source and confusing AU the following conditions:

*<u>Condition 1</u>*: the Co-Information between AU intensity, MEG_t_ activity and the confused categorizations–i.e., comparing single-trial “disgust” or “anger” categorizations vs. other categorizations:

Co-I = MI(AU intensity; MEG_t_) + MI(confused responses; MEG_t_) - MI(AU intensity, confused responses; MEG_t_)

*<u>Condition 2</u>*: the Co-Information between AU intensity, MEG_t_ activity and the discriminative categorizations, i.e., single-trial “disgust” vs. “anger” categorizations:

Co-I = MI(AU intensity; MEG_t_) + MI(discriminative responses; MEG_t_) - MI(AU intensity, discriminative responses; MEG_t_)

On trials showing disgust and anger stimuli and presenting both the confusing and discriminative AUs, we computed for each source and discriminative AU the following conditions:

*Condition 1*: the Co-Information between AU intensity, MEG_t_ activity and the confused categorizations.

*Condition 2*: the Co-Information between AU intensity, MEG_t_ activity and the discriminative categorizations.

For each of these 4 computations, we repeated for 200 times Co-I computations with shuffled MEG_t_, using as statistical threshold the 95th percentile of the distribution of 200 maxima (each taken across the source x time Co-I matrix of each shuffled repetition).

For each participant and AU, we performed independent-sample t-tests on the two Co-I (i.e., with confused vs. discriminative responses) at AU representation peak–i.e., MI(AU intensity; MEG_t_) peak. Figure 6 shows the group-mean significant Co-I for confusing AUs and discriminative AUs.

## Acknowledgements

This work was funded by the Wellcome Trust (Senior Investigator Award, UK; 107802) and the Multidisciplinary University Research Initiative/Engineering and Physical Sciences Research Council (USA, UK; 172046-01), awarded to Philippe G. Schyns; ERC [FACESYNTAX; 759796], awarded to Rachael E. Jack; and the Wellcome Trust [214120/Z/18/Z], awarded to Robin A.A. Ince. The funders had no role in study design, data collection and analysis, decision to publish or preparation of the manuscript.

## Supplemental materials

**Supplemental Figure S1.**
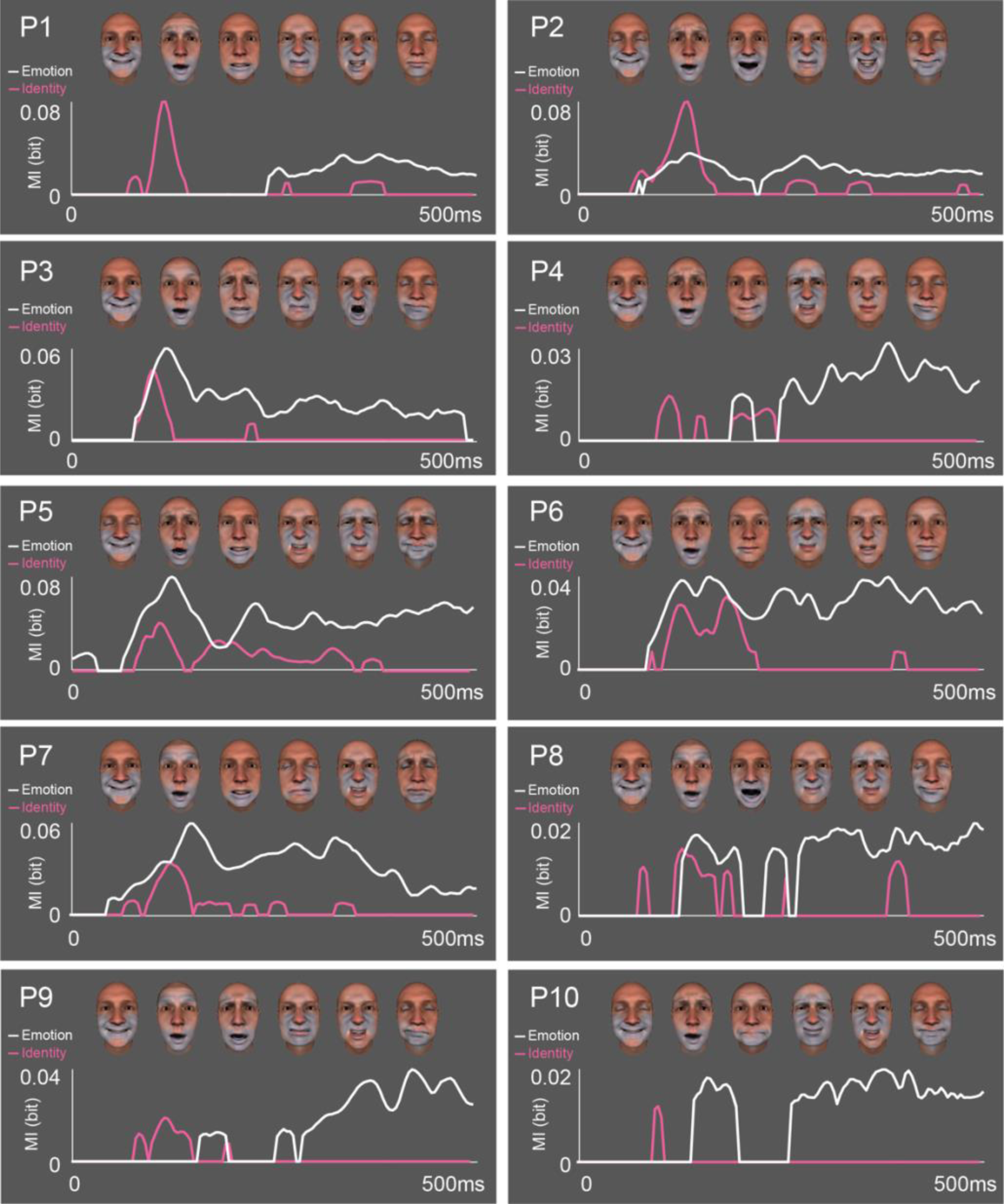
Dynamic representations of categories (individual participant data). Emotions. For each participant, we show their emotion models (from left to right: happy, surprise, fear, disgust, anger, sad). We computed the dynamic representations of the emotion models as MI(MEG_t_; 6 emotion category) and identity representation MI(MEG_t_; 8 identities), on all sources of the three pathways (see Table 1), every 4ms between 0 and 500ms post animation onset (FWER-corrected over 2,775 sources × 125 time points, *p*<0.05). The white curve shows the maximum emotion representation across all sources. **Identities.** As a control, we repeated this representation analysis using task-irrelevant face identity. We computed MI(MEG_t_; face identity label), every 4ms between 0 and 500ms from face onset (FWER-corrected over 2,775 sources × 125 time points, *p*<0.05). The magenta curve shows the maximum MI across sources. The white (vs. pink) curve shows the cross-source average of the maximum emotion (vs. identity) representation across all sources.

**Supplemental Figure S2.**
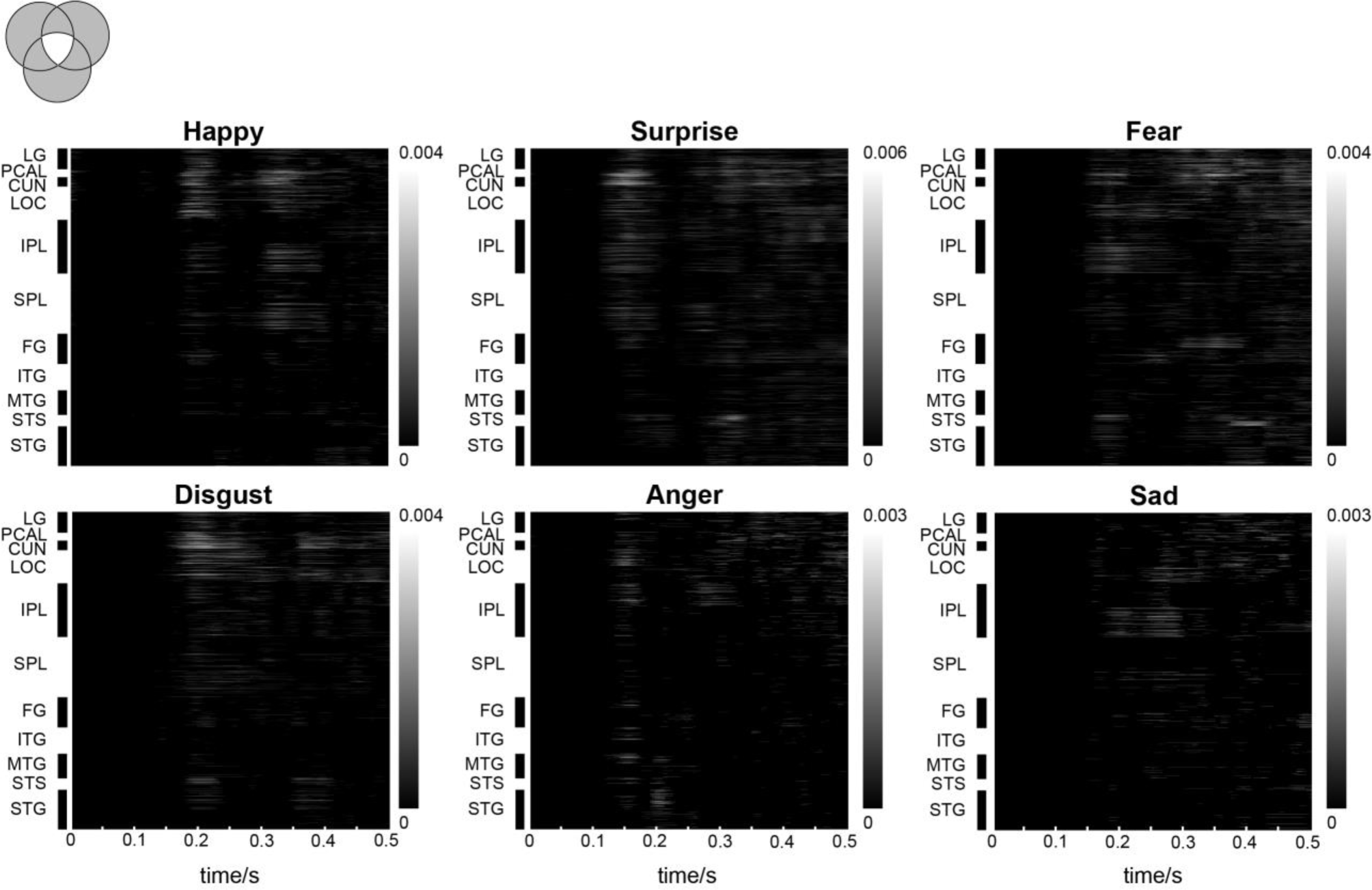
Emotion representations. For each emotion category, we traced the dynamics of its representations across the brain, computing MI(MEG_t_; responding to this emotion vs. “others”) on all sources of the three pathways (see Table 1), every 4ms between 0 and 500ms from animation onset (FWER-corrected over 2,775 sources × 125 time points, *p*<0.05). The matrices show the averaged source (Y-axis) × time (X-axis) MI representations across all participants.

**Supplemental Figure S3.**
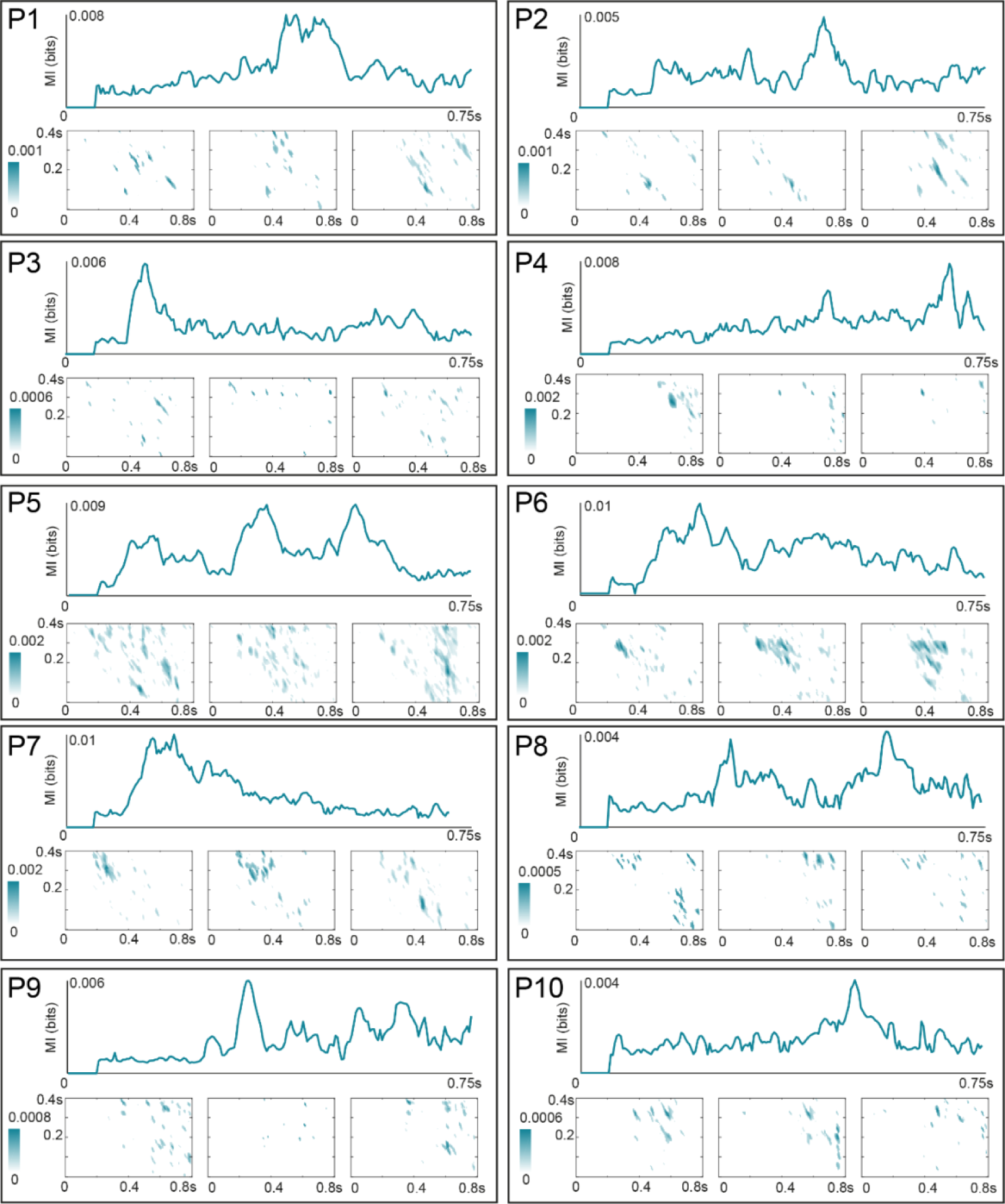
Individual AU representations and communications. For each participant per panel, we computed the dynamic representations of each AU into MEG activity as MI(MEG_t_; AU intensity), FWER-corrected over 2775 sources × 200 time points, *p*<0.05. <u>Curve</u> shows the maximum MI representation across ventral, dorsal and social pathways sources. For each AU, in each emotion model, we selected the sender (in occipital cortex) and the receiver (independently for MTG, STS and STG) with maximum significant MI. For each AU, we computed DFI communications between sender and receiver sources every 4 ms between 0 and 600ms post animation onset in the receiver time course (x-axis), and for every 4ms delay between 0 and 400ms from the sender (y-axis). <u>Matrix plot</u> reports per-participant DFI communications to MTG (left), STS (middle) and STG (right) as the mean of significant DFI across all AUs–FWER, *p*<0.05.

**Supplemental Figure S4.**
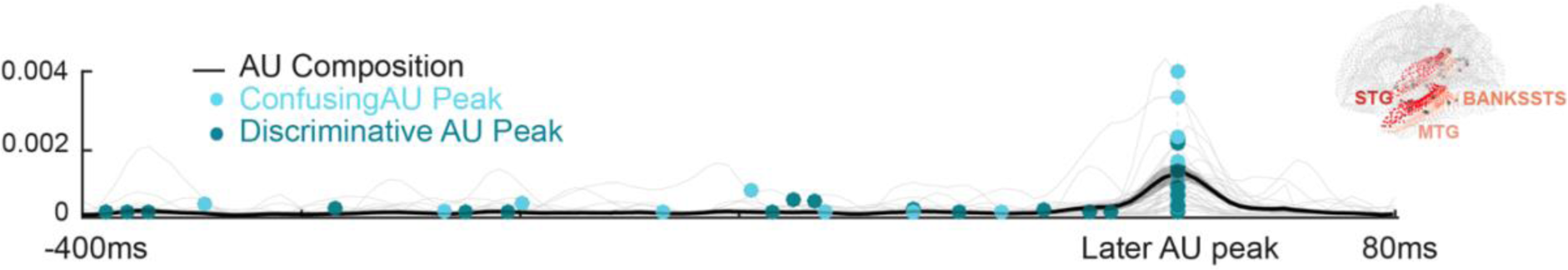
For each participant, we quantified in all MTG, STS and STG sources the composition of each confusing AU with each discriminative AU, computing Synergy(confusing AU intensity; discriminative AU intensity; MEG_t_). The black curve shows the cross-participant mean of maximum composition across all sources, aligned at peak representation time of the latest AU. For each AU pair, confusing (light blue) and discriminative (dark blue) AUs, we plotted the curve of their synergistic composition (thin grey curves). Our results reveal that AU compositions occur after both confusing and discriminative AUs converge on MTG, STS or STG sources.

